# Mechanism of Voltage Gating in the Voltage-Sensing Phosphatase Ci-VSP

**DOI:** 10.1101/2022.02.17.480971

**Authors:** Rong Shen, Yilin Meng, Benoît Roux, Eduardo Perozo

## Abstract

The conformational changes in voltage-sensing domain (VSD) are driven by the transmembrane electric field acting on charges and countercharges. Yet, the overall energetics and detailed mechanism of this process are not fully understood. Here, we determined free energy and displacement charge landscapes, as well as major conformations corresponding to a complete functional gating cycle in the isolated voltage-sensing domain of the phosphatase Ci-VSP (Ci-VSD) comprising four transmembrane helices (segments S1-S4). Molecular dynamics simulations highlight the extent of S4 movements. In addition to the crystallographically determined activated ‘Up’ and resting ‘Down’ states, the simulations predict two novel Ci-VSD conformations: a deeper resting state (‘Down-minus’) and an extended activated (‘Up-plus’) state. These additional conformations were experimentally probed via systematic cysteine mutagenesis with metal-ion bridges and the engineering of proton conducting mutants at hyperpolarizing voltages. These results show that voltage activation involves sequentially populating these four states in a stepwise way, translating one arginine across the membrane electric field per step, transferring ~3 e_0_ charges.

## Introduction

Voltage-sensing domains are conserved functional modules that transduce voltage-driven charge rearrangements to exert conformational biases on effector domains present in a variety of membrane proteins^1–4^, including ion channels, solute carriers and phosphatases. They form an antiparallel bundle of four transmembrane helices (segments S1-S4). A series of positively charged high-pKa residues (mostly arginines) located on the S4 segment act as the key structural element for voltage sensing^5^. The S4 gating charge motif varies in different VSDs^6,7^, a fact that leads to a diversity of voltage sensing profiles. These positive charges make extensive and sequential salt bridge interactions with negative countercharges on segments S1-S3 (and lipid headgroups) upon S4 translocation^8–11^. At the center of VSDs, a single layer of in-plane hydrophobic residues on the S1-S3 segments, referred to as the hydrophobic plug, the charge-transfer center or the hydrophobic gasket^12–15^ electrically separates the intra- and extracellular facing water-permeable VSD vestibules, leading to a dramatic focusing of the electric field^16,17^.

In addition to mediating ion and/or proton conduction across the membrane, VSDs are able to precisely control enzyme activity in voltage-sensing phosphatases. Ci-VSP was the first member of the VSP family that showed robust membrane potential dependent enzymatic activity towards inositol phospholipids, a type of crucial secondary messenger lipid responsible for many basic cellular functions^4,18^. Ci-VSP has a N-terminal transmembrane VSD and a C-terminal cytosolic phosphatase domain, homologous with the tumor suppressor phosphatase PTEN (protein lipid phosphatase and tensin homolog deleted on chromosome 10), but with a broader substrate specificity than PTEN^19–23^. Ci-VSP is likely a monomer in the native membrane environment and functions independently even at high expression level^24,25^. Furthermore, a single mutation at the catalytic center of the phosphatase domain of Ci-VSP can eliminate its enzymatic activity with little effect on electrical sensitivity of the voltage-sensing domain^4^, making Ci-VSP an ideal model for studying the gating mechanism of the VSD itself.

To date, there is no atomic structure of any full-length VSPs available, however crystal structures of Ci-VSD have been determined in two different states (‘Up’ and ‘Down’, Extended Data Fig. 1a), providing an initial atomistic view of the voltage-driven conformational change within a given VSD^12^. A recent set of cryo-electron microscopy (cryo-EM) structures of voltage-dependent ion channels has confirmed and greatly expanded these initial results^26–29^. From the two Ci-VSD structures, the S4 movement is best characterized by a helix turn and axial translocation (three-residue; or one ‘click’) movement, while S1-S3 display little or no conformational changes. This movement is broadly consistent with the classic helical screw model, whereby the S4 helix translates along and rotates around its main axis^30,31^, resulting in the translocation of one arginine residue across a region near the center of the VSD where the transmembrane electric field is focused to yield an estimated gating charge of ~1 *e*_0_^12^. However, voltage clamp fluorometry and enzyme activity assays show that Ci-VSP has distinct substrate preferences at different depolarization membrane potentials, pointing to the likelihood of multiple Ci-VSD conformationally stable states undergoing multistep conformational transitions during a complete functional cycle^20,23,24^. Here, we evaluate the full conformational space of a VSD as a function of voltage and displacement by combining: 1- The computation of free energy landscapes in Ci-VSD; 2- Estimating the number of energetically stable conformations and 3- Experimentally evaluating them by means of metal-ion bridges and the engineering of proton conducting mutants. This interrelated approach offers the most detailed insight to date on the mechanism of electric field transduction in VSDs.

## Results and Discussion

### Free energy landscapes and major functional states governing Ci-VSD gating cycle

Inspired by the charge-countercharge ‘click’ model proposed from the structural study of Ci-VSD^12^, we evaluated the existence of extended S4 movements beyond those observed experimentally. This hypothesis is based on the assumption that additional charge-countercharge partnerships are possible with additional S4 movements and limited changes in the S1-S3 countercharges. Any additional S4 movements beyond the experimentally determined Up and Down conformations might lead to the identification of novel, energetically favorable charge-countercharge interactions. To this end, we generated two homology models of Ci-VSD in the ‘Up-plus’ (one ‘click” beyond Up) and ‘down-minus’ (one ‘click’ below Down) states, and equilibrated them using molecular dynamics (MD) simulations (Extended Data Fig. 1a-c). To characterize the stability of these states and inform the details of the gating mechanism, we calculated the free energy landscapes governing the conformational transition process and the related displacement charges, *Q*_d_ = 〈∑*_i_ q_i_z_i_*〉, representing the coupling between the transmembrane potential and the VSD (Fig. 1a, b). Observable gating charges are determined from the difference in *Q*_d_ between two conformations. Two-dimensional potentials of mean force (2D-PMFs) at three membrane potentials (0 and ±150 mV) were calculated using the Hamiltonian-replica exchange molecular dynamics (H-REMD) umbrella sampling simulations (Extended Data Fig. 1d-g), with the translocation and rotation of S4 being used as reaction coordinates to describe the conformational change.

**Figure 1.**
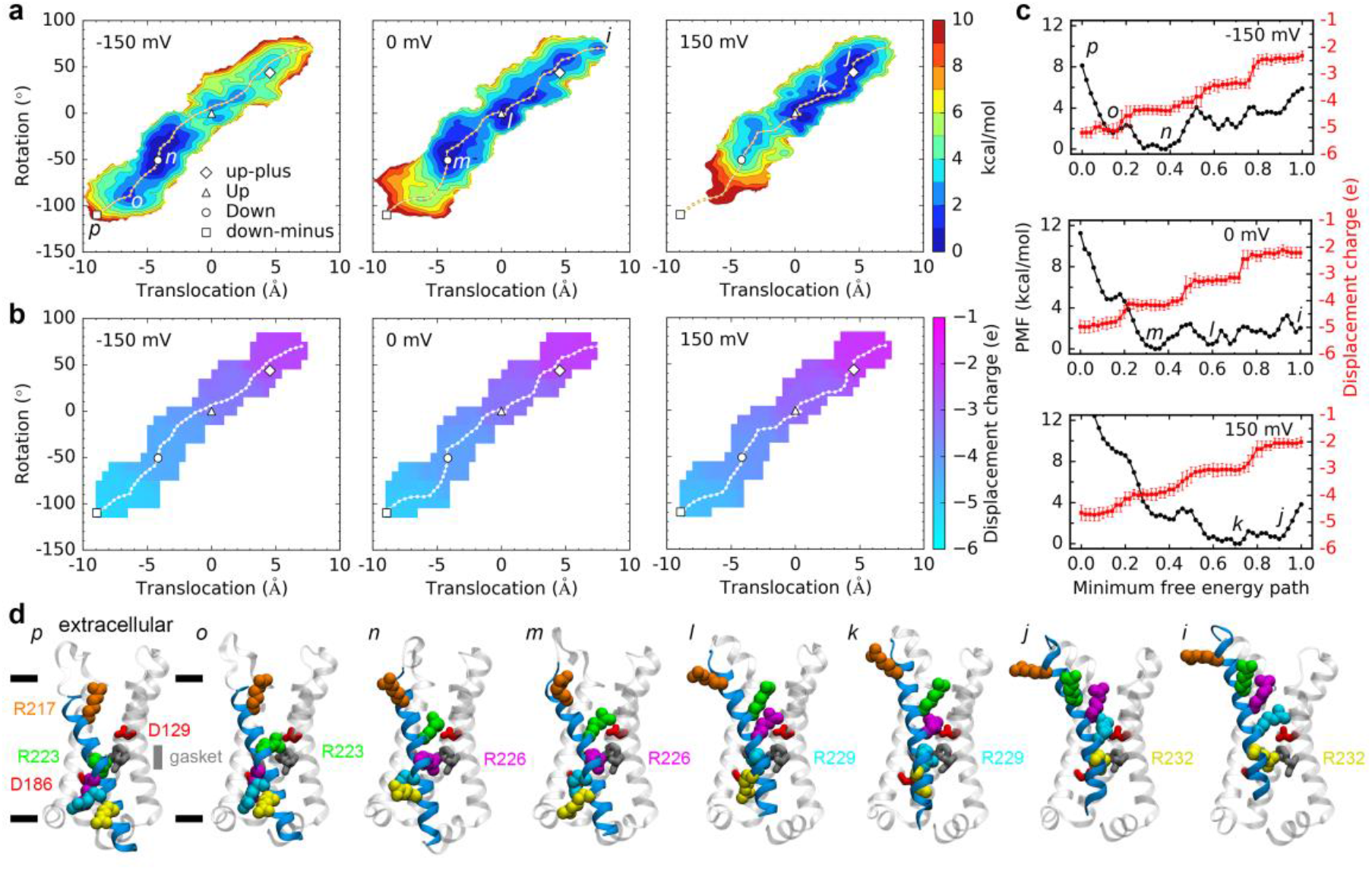
Free energy landscapes and gating charges in the movement of the voltage-sensing domain S4 segment at different membrane potentials. **a**, Two-dimensional PMFs of the gating process at −150 mV, 0 mV and 150 mV. Translocation and rotation of the S4 segment along and around its principal axis are used as the reaction coordinates, respectively. The dashed lines represent the minimum free energy paths (MFEPs). Projection of the four starting configurations onto the two reaction coordinates and representative local minima along the MFEPs are highlighted. **b**, Average displacement charge, *Q*_d_ = 〈∑*_i_q_i_z_i_*〉, at different membrane potentials. Snapshots (*n* = 50) from the last 5 ns trajectories of the H-REMD simulations were used for the calculation. The dashed lines are the MFEPs as in **a**. **c**, One-dimensional PMF and the average displacement charge along the MFEP at different membrane potentials. Error bars denote standard deviation. **d**, Snapshots of the representative local minima. The positive charged residues on the S4 segment are shown in the van der Waals (vdW) representation with different colors. The two countercharges D129 (S1) and D186 (S3), and the hydrophobic gasket forming residues I126 (S1), F161 (S2) and I190 (S3) are displayed as sticks. The gray and black bars represent the approximate position of the hydrophobic gasket and membrane bilayer, respectively.

From these calculations the membrane potential biased the free energy landscape in the expected directions: a negative potential increased the population of the ‘down-minus’ state, a positive potential increased the population of the ‘Up-plus’ state, and the ‘Down’ and ‘Up’ states have the lowest free energies at 0 mV as anticipated from the crystallographic data. Interestingly, the ‘Up-plus’, ‘Up’ and ‘Down’ states were quite stable, populating at local free energy minima, at all three membrane potentials. No so in the case of the ‘down-minus’ state, which was found to be energetically unfavorable compared with the other three states, even if it stayed near its starting conformation during the initial MD simulation (Fig. 1a, Extended Data Fig. 1c). Distinctive free energy basins corresponding to intermediate metastable states have also been observed in the free energy landscapes (Fig. 1a, c). Analysis of the trajectories showed that these metastable states represent an interval backbone movement of S4 compared with the adjacent main states. These movements take place while keeping the key salt bridge interactions between the side-chains of the gating residues on S4 and the countercharge residues on S1-S3 or lipids mostly unchanged (Fig. 1d). This result suggests that the S4 backbone movement likely takes place ahead of the rearrangement of the side chain salt bridge interactions during gating. This is also in agreement with the displacement charge calculation result showing that only the conformational change between two adjacent main states contributes to an estimated gating charge of ~1 *e*_0_ by moving an arginine across the electric filed focused at the center hydrophobic gasket (Fig. 1b, c). Such that a whole functional cycle movement from the ‘down-minus’ state to the ‘Up-plus’ state contributes a total gating charge of ~3 e_0_..

### A general ‘click’ model of VSD rearrangements

The secondary structure of S4 (α-helical and/or 3_10_-helical) has attracted a lot of attention and debate, as it is directly related to the orientation and interacting network of the gating residues in different states, and thus the detailed VSD gating mechanism has converged towards two mechanistically related models: A “helical screw model”, where the S4 has both rotational and translational movement or a “helical sliding model” where the S4 translates along its principal axis with minimal rotational movements^26,29^. Both, crystal structures and MD trajectories along the minimum free energy path of one of the free energy landscapes, reveal that the Ci-VSD S4 segment tends to adopt a fully α-helical conformation during gating with only one or two residues occasionally adopting a 3_10_-helical conformation. This high α-helical propensity was not a result of the harmonic restraints applied in the PMF calculations (Extended Data Fig. 2a-c). We suggested that it is the location of the countercharges (in other words the hydrogen-bonding network), what shapes the secondary structure of S4. To test this hypothesis, MD simulations were performed with a ‘Up-plus’ state model of Ci-VSD with double mutant D129S/A154E, in reference to the position of countercharges in the VSDs of potassium channels. During a 100 ns production MD run, the S4 segment adopted a perfect α-helical conformation. However, after two hydrogen-bonding interactions between the gating residues and countercharges were enforced at the very beginning of the simulation, the extracellular region of S4 spontaneously changed into a 3_10_-helix in the following simulation without any restraints. This α-helix to 3_10_-helix transition placed the four arginine residues all toward the aqueous crevice of Ci-VSD and increased the hydrogen-bonding probability of the gating residues with the countercharges (Extended Data Fig. 2d-h).

Given recent breakthroughs in cryo-EM, a number of VSD structures in two different states have now been determined. These enable us to inspect the movement of other VSDs in an alternative way using the experience gained from the study of Ci-VSD. The S4 segment of the VSDs with available structures has three major secondary structure conformations: α-helical^12^, 3_10_-helical^26^ and a combination of α-helical and 3_10_-helical^29^. This pattern is maintained throughout different states of the same VSD although S4 is not always a single continuous helix, e.g., in the HCN channel, the S4 segment breaks into two helices in the hyperpolarization state (Extended Data Fig. 3a-c). Superimposition of the two structures of the same VSDs shows that the conformation of the S1-S3 segments is quite stable, and the overall position of gating residues and the region of the α-helix/3_10_-helix along S4 in relative to S1-S3 remain approximately the same, as shown in the MD simulation study of the Shaker K^+^ channel VSD^10^. It implies that one gating residue will occupy its adjacent gating residue’s position in a one-step movement of S4, just as what was observed experimentally in Ci-VSD. For the S4 segment having both α-helical and 3_10_-helical regions, a continuous α-helical to 3_10_-helical or 3_10_-helical to α-helical transition will take place for residues in the boundary of the two helical conformations. To fulfill this type of movement, the α-helical S4 segment will translate along and rotate around its axis (the conventional helical screw motion), the 3_10_-helical S4 segment will translate along its axis without any rotation (the conventional helical sliding motion), and the α-helical/3_10_-helical S4 segment will undergo both translocation and rotation at the α-helix region and translocation alone at the 3_10_-helix region (a combination of helical screw & sliding motion).

Thus, the sequential stepwise movement of S4 can be simplified and generally described with a set of discrete ‘clicks’, so that one helix turn or one ‘click’ represents the basic element of the movement of S4, and corresponds to a complete transition between two adjacent major states. The translocation and/or rotation motion of S4 is related to the secondary structure of S4, which is thought to be regulated by the location of the countercharges and the requirement of shielding the electrostatic destabilization of the gating residues^29^. Using the KAT1 potassium channel as an example^6^, if the central region of S4 does not adopt a 3_10_-helical conformation, the arginine residues on the extracellular side will face the hydrophobic VSD-pore domain interface or the hydrophobic lipid tails, which is highly energetically unfavorable (Extended Data Fig. 3d-g).

### State dependent residue-residue interactions probed by Cd^2+^ bridges

Efforts to obtain structural correlates of the two predicted states, ‘Up-plus’ and ‘down-minus’, have been frustrated by the size, biochemical stability and/or crystallization hurdles. However, we have carried out a metal bridge experimental strategy to evaluate the extended conformational landscape in Ci-VSD, characterize all predicted conformations and explore the existence of other putative states. To this end, we introduced cysteine mutations in the VSD, guided by our computational models, and evaluated the formation of Cd^2+^ bridges to identify the dynamic residue-residue interactions during the functional cycle^10,13^. Combined with restrained MD simulations with an explicit Cys-Cd^2+^-Cys bridge^32,33^, we then established the most likely state at which each specific interaction can be formed.

We studied the Cd^2+^ effect on the net OFF gating charge (*Q*) of single cysteine mutants using the cut-open oocyte voltage clamp technique (Extended Data Fig. 4). For wild type (WT) Ci-VSD, single S1 (M133C, L137C), S2 (A154C) and S3 (T197C) and the intracellular side of S4 (L224C-R232C) mutants, 100 uM Cd^2+^ has little effect on the normalized *Q* at the maximum depolarization potential (Fig. 2a). However, the *Q-V* curve of A154C shifted towards more positive voltages compared to others. This effect can be restored after washing with DTT or the metal chelator EGTA (Fig. 2b). In contrast, A154E showed little response to the external Cd^2+^ over the whole range of the voltage pulses. We interpreted this result to suggest that Cd^2+^ could get into the external crevice and bind at the site near A154C, slowing the movement of S4 at lower voltages as shown in the gating current traces (Extended Data Fig. 4a, b). It is worth noting that the mutant R223C enabled a continuous proton current through the VSD at negative potentials, consistent with observations in the mutant R223H^34^ and analogous mutants of ion channels^35,36^. Thus, changes in steady state currents were used in the study of the Cd^2+^ effect on R223C (Extended Data Fig. 4c). Surprisingly, an apparent decrease of the maximum charge was observed for Y200C. As there are plenty of negative residues at the extracellular face of the VSD, potential formation of Cd^2+^ bridges between the cysteine and glutamates/aspartates could be responsible for the net charge reduction^10^. We found that neutralization of D136 (on S1) and each of the three negative residues on the S3-S4 loop (D204, E205 and E209) can lead to a partial relief of the Cd^2+^ effect, while a quadruple countercharge mutant (D136A/D204A/E205A/E209A) almost totally eliminated the Cd^2+^ effect and increased the kinetics of gating charge movement dramatically (Extended Data Fig. 5a-c). MD simulations of Ci-VSD/Y200C showed that the Cd^2+^ binding at Y200C is located along the gating charge transfer pathway. So that this bridge could impede the movement of the arginine residues on S4 and thus decrease the net charge as in A154C. In addition, the S3-S4 loop was found to be very flexible in our MD trajectories, hence the Cd^2+^ bridge between Y200C and the S3-S4 loop could restrict the rearrangement of the loop, and in turn the movement of S4 (Extended Data Fig. 5d, e).

**Figure 2.**
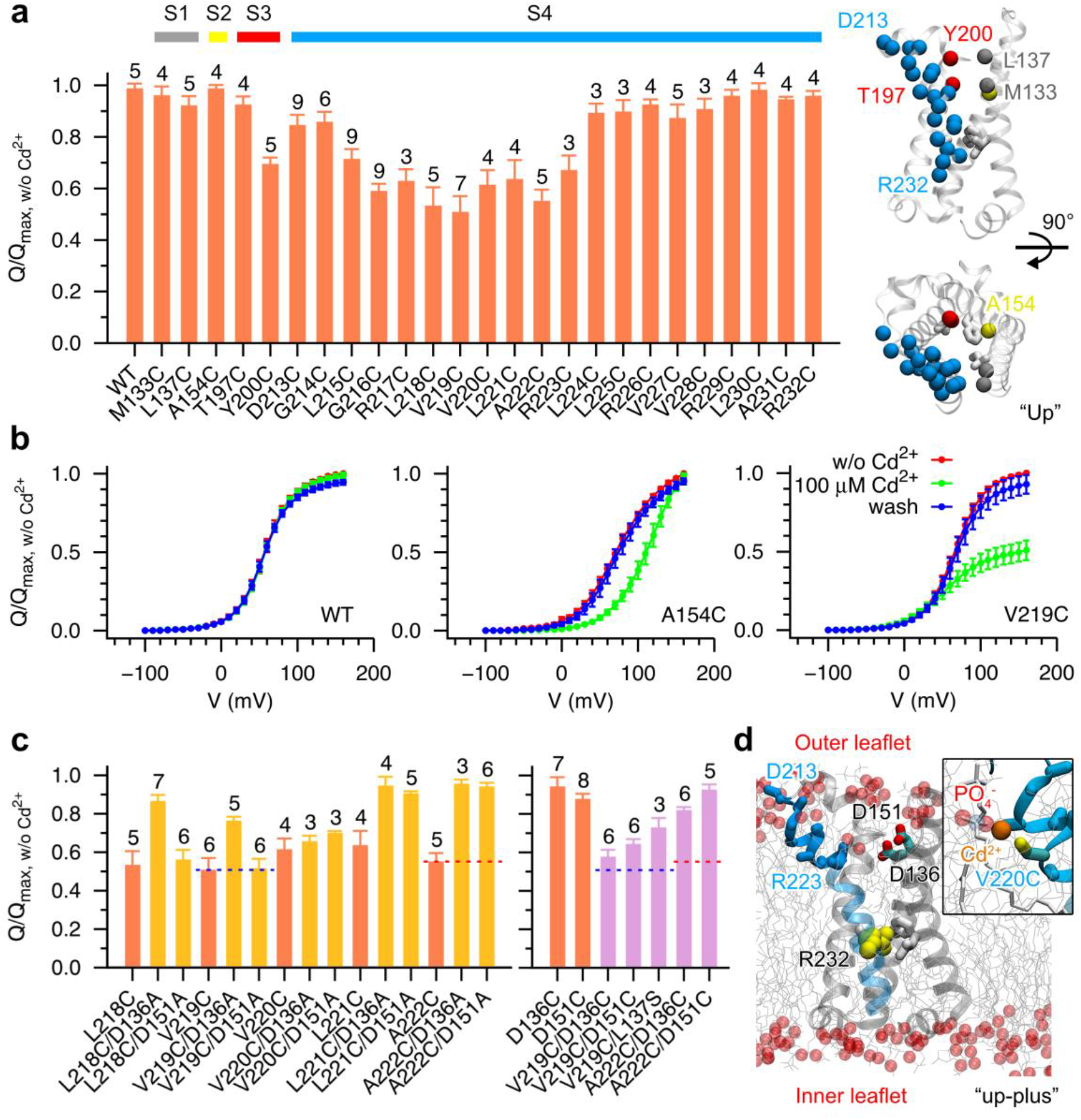
Effect of extracellular cadmium (Cd^2+^) ions on Ci-VSD charge movements from single cysteine mutations. **a**, Left, the relative immobilization of OFF gating charge movement of wild type (WT) and single cysteine mutants of Ci-VSP at the maximum voltage in response to 100 μM Cd^2+^. The net charge *Q* is calculated by numerically integrating the OFF gating current, except for the R223C mutant that the maximum current at the end of the voltage pulse was used for the normalization and comparison. Error bars denote standard deviation with the number of experiments being listed on the top of each of the bars. Right, side and top view of the “Up” state VSD (PDB: 4G7V) with carbon alpha atoms of the mutants being highlighted in spheres with different colors: gray (on S1), yellow (on S2), red (on S3) and blue (on S4), and the hydrophobic gasket forming residues in white sticks. **b**, Representative charge-voltage (*Q*-*V*) curves in the absence (red), in the presence (green) and after washout (blue) of 100 μM Cd^2+^. The *Q-V* curves were normalized with respect to the maximum net charge without (w/o) Cd^2+^ for each experiment. Error bars denote standard deviation (WT, *n* = 5; A154C, *n* = 4; V219C, *n* = 7). **c**, Countercharges D136 and D151 play indirect roles on external Cd^2+^ induced immobilization of OFF gating charge movement of single cysteine mutants on the extracellular side of S4 other than forming direct metal bridges. **d**, A snapshot of the “Up-plus” state VSD from molecular dynamics simulations. Only parts of the lipid molecules (gray lines) and their non-ester phosphate oxygen atoms (red spheres) are shown for clarity. The extracellular (residues: 213–223) and intracellular sides of the S4 helix are shown in solid and transparent blue ribbon representation, respectively. The alpha and beta carbon atoms of these extracellular side residues are highlighted in ball-and-stick representation. Insert is an illustration of a potential Cd^2+^ bridge between a cysteine on S4 and a non-ester phosphate oxygen atom from the outer leaflet.

To identify the factors underlying the Cd^2+^ induced net charge reduction of the single cysteine mutants on the extracellular side of S4 (D213 to R223), we made additional mutants especially of the two negative residues D136 and D151, as they may form endogenous Cd^2+^ bridges with the cysteines. For L218C and V219C, the D136A mutation decreased the Cd^2+^ effect, but not for D151A. For V220C, both mutations showed negligible effect. While, for L221C and A222C, either D136A or D151A could rescue the Cd^2+^ effect, making the results inconclusive (Fig. 2c). To further test whether the single mutants would make Cd^2+^ bridges with the aspartates, we mutated them to a cysteine. For V219C and A222C, an additional cysteine mutation at position 136 or 151 both decreased the Cd^2+^ effect in contrast to the results of the single cysteine mutants, eliminating the possibility of formation of a Cys-Cd^2+^-Cys bridge. In addition, a negative control mutation, L137S, can also relieve the Cd^2+^ effect.

No periodicity was detected in the reduction of the maximum net charge and the shift of the half-maximum activation voltage (*V*_1/2_) (Extended Data Fig. 6a), pointing to factors other than the Cys-Cd^2+^-Asp bridge likely underlie the Cd^2+^ effect. We then evaluated the possibility that these effects were mediated by Cys-Cd^2+^-lipid interactions. We increased the external Cd^2+^ concentration to 1 mM, and found a notable decrease of the net charge for A222C/D136A and A222C/D151A, but not for L225C/D136A and L225C/D151A which were used as a control, considering the concentration dependent Cd^2+^ effect on the single mutants (Extended Data Fig. 6b-e). Using V219 as an example, analysis of the MD trajectories suggested that the residues on the extracellular side of S4, are not in a favorable distance to form Cd^2+^ bridges with D136 and D151. However, they are positioned to make favorable contacts with the outer leaflet of lipids during S4 movement. Interestingly, this putative contact stopped right at R223 at the ‘Up-plus’ state (Extended Data Fig. 6f-g). The agreement between the experimental and computational results suggest that specific single cysteine mutants can form Cd^2+^ bridges with the phosphate oxygen atoms of membrane phospholipids, a result that is fully consistent with the ‘Up-plus’ state as the outermost state of Ci-VSD (Fig. 2d).

Based on results from the single Cys mutants, we evaluated the Cd^2+^ effect on double mutants to investigate the residue-residue interactions that might define further structural constraints between S4 and S1-S3. The T197C on S3 was first examined against the cysteine mutants on S4 (Fig. 3a). By taking into account the effect of single mutants, we found four notable bridging sites with a periodicity of three residues overlapping with the positions R217C, V220C, R223C and R226C (Fig. 3b). To fully explore this conformational space, we also examined the partner mutants on S1 (M133C and L137C) and S2 (A154E), and found nine additional bridges (Fig. 3a, b). As in the single Cys mutants experiments, the reduced gating currents of double mutants could be restored after the wash out (Fig. 3c). Restrained MD simulations of the double mutants forming Cd^2+^ bridges were then carried out to verify the most likely state(s) at which each bridge forms (Fig. 3d, Extended Data Fig. 7a-d). In short, we built models for each double mutant at all the four states: “down-minus”, “Down”, “Up” and “Up-plus”. By analyzing the distance distribution of the beta carbon atoms of the two mutated residues, we were able to evaluate the bridge forming state. An explicit Cd^2+^ bridge was then introduced into the specific model by inserting a Cd^2+^ ion and protonating the cysteine residue(s)^37^. We note that because of the lack of accurate force fields for the metal bridge, external bond and angle restraints needed to be imposed to maintain the Cd^2+^ bridge^33^. However, slight movements of S4 can always achieve an appropriate bridge while avoiding the breakdown of its secondary structure as shown by the backbone root-mean-square deviation (RMSD) of S4. These bridges can be perfectly projected onto their respective structures (Fig. 3e). Together with information from the unpaired double mutants, we conclude that other than the four major states evaluated here, it is unlikely that the S4 segment is able to populate additional conformational states under physiological conditions. For instance, R217C can form a bridge with T197C in the ‘down-minus’ state, but the one helix turn above residue G214C is clearly unable to form stable interactions with T197C. Also, D213C and G214C can form bridges with L137C in the ‘down-minus’ state but are unable to form bridges with M133C which is one helix turn below L137C. These results lead us to conclude that S4 cannot move further down from the ‘down-minus’ state (itself a low probability event). For the upward S4 movement, R229C/A154E and R226C/M133C can form bridges at the ‘Up-plus’ state, but R232C/A154E and R229C/M133C cannot form bridges, implying that the ‘ Up-plus’ state is the terminal state in any activating upward gating movement of S4. Based on restrained MD simulations we also found that distance should not be the only criteria that determines successful Cd^2+^ bridge formation. The space between two cysteine residues may have been previously occupied by other bulky or charged residue(s) preventing Cd^2+^ binding, or side chain rotamer variability could lead to unfavorable orientations while still in a bridging distance, inhibiting bridge formation without unfolding the helix (Extended Data Fig. 7e-h). Therefore, interpretation and design of metal bridges in the absence of structural information must be done with a healthy dose of caution.

**Figure 3.**
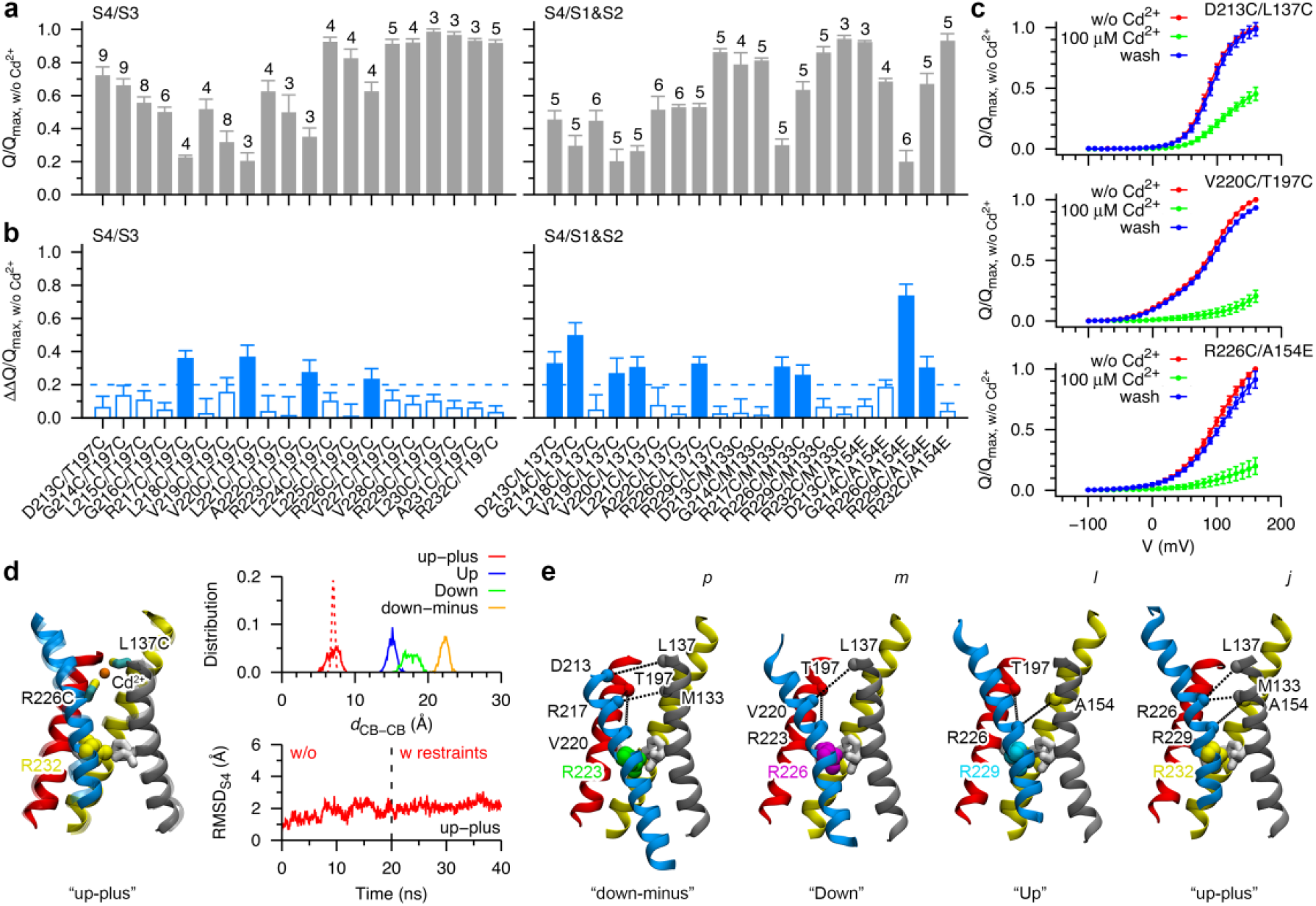
State dependent Cd^2+^ bridges formed by residues on the S4 and S1-S3 helices. **a**, The relative immobilization of OFF gating charge movement of double mutants of Ci-VSP at the maximum voltage in response to 100 μM Cd^2+^. Error bars denote standard deviation with the number of experiments being listed on the top of each bar. For mutants including R223C, the current amplitude at the end of the voltage pulse was used for the normalization and comparison. **b**, The effective reduction of the normalized maximum net charge of the double mutants in **a** by considering the Cd^2+^ effect on each single mutant. A threshold of 0.2 (dashed line) was used to highlight the most likely interacting pairs (solid bar). Error bars denote standard deviation. **c**, Representative normalized *Q*-*V* curves of double mutants in the absence (red), in the presence (green) and after washout (blue) of 100 μM Cd^2+^. Error bars denote standard deviation with the same sample sizes as in **a**. **d**, Left, comparison of the conformation of the “Up-plus” state Ci-VSD before (transparent) and after (solid) a restrained MD simulation with the R226C-Cd^2+^-L137C bridge. Right, distance distributions (*n* = 1,000) of the beta carbon atoms of R226C and L137C in the MD simulations of the double mutant Ci-VSD (R226C/L137C) in different states (solid line) and in the restrained MD simulation of the double mutant Ci-VSD in the “Up-plus” state only (dashed line) (top), and the root-mean-square deviation (RMSD) of the backbone atoms of S4 in the MD (black line) and restrained MD (red line) simulations of the “ Up-plus” state Ci-VSD mutant (bottom). During the restrained MD simulation, the geometry of the Cys-Cd^2+^-Cys bridge was harmonically restrained. **e**, Projection of the state-depended Cd^2+^ bridge interactions suggested by restrained MD simulations into the four conformational states of Ci-VSD: “down-minus” (D213C/L137C, R217C/M133C, R217C/T197C and V220C/T197C), “Down” (R223C/T197C and V220C/L137C), “Up” (R226C/T197C and R226C/A154E) and “Up-plus” (R226C/L137C, R226C/M133C and R229C/A154E). The alpha carbon atoms of interacting residues are shown in spheres and connected with dashed lines. The hydrophobic gasket forming residues and the arginine residue in the middle of the gasket are shown in sticks and vdW representation, respectively.

### State dependent proton currents in single Ci-VSD S4 mutants

Single mutations of gating charge arginines on S4 have been found to cause ionic leak through the VSDs of voltage-gated ion channels^34–36,38,39^, the so-called ‘omega current’ or ‘gating pore current’. Continuous inward currents were also observed in our recording of R223C at potentials more negative than −20 mV. As *N*-methyl-D-glucamine (NMDG) was the only cation in our external recording buffer, we suspected that the inward currents were carried by protons. To test whether protons can permeate through other single mutants, we recorded the gating currents again at different external pH values (pH_o_). A clear effect was observed on the mutants of three gating charge arginines (R223C, R226C and R232C), especially the two outermost gating residues, in agreement with the study of single histidine substitution of these residues^34^. Other S4 single cysteine mutants had minimal effects on the net charge, although the kinetics were faster at high pH_o_ than at low pH_o_ (Fig. 4a, Extended Data Fig. 8a, b). In the case of both, R223C and R226C current magnitude increased at lower pH_o_, yet each mutant showed distinct current characteristics (Fig. 4b). R223C facilitated a steady leaky proton current similar to the typical ‘omega current’, while R226C exhibited a gating current like transient leaky proton current. This suggests that proton leaks through R226C only at certain intermediate state(s) before the VSD finally reaches its terminal “down-minus” state, a state incompatible with R226C-carried omega currents.

**Figure 4.**
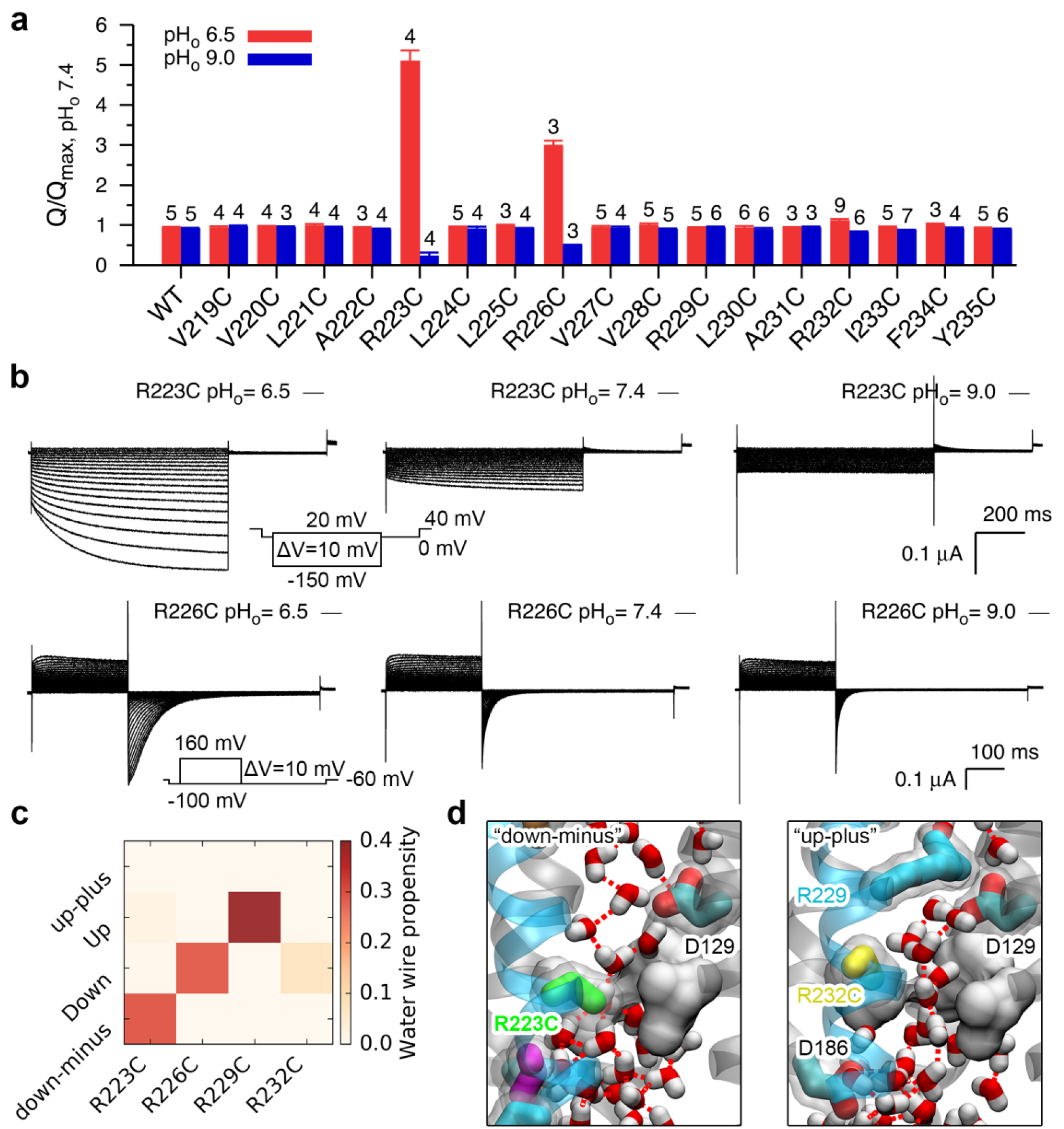
State dependent proton conduction through single cysteine mutants informs on the Ci-VSD conformations. **a**, Changes of the maximum OFF net charge of WT and single cysteine mutants of Ci-VSP in response to the acidification of the external buffer. The net charge *Q* was normalized with respect to the maximum value at pH_o_ 7.4. The current amplitude of R223C at the end of the voltage pulse was used for the normalization and comparison. Error bars denote standard deviation with the number of experiments being listed on the top of the bars. **b**, Representative currents for R223C and R226C with the external recording buffer at different pH values. Insets are the voltage pulse protocols used to elicit the currents. **c**, Propensity of the formation of a continues hydrogen-bonded water wire connecting both sides of the mutated VSD at different states. Snapshots from the last 30 ns trajectory of each 50 ns MD simulation were used for the calculation (*n* = 3,000). **d**, Snapshots of the R223C and R232C mutants at the “down-minus” and “Up-plus” states, respectively. Instantaneous hydrogen bonds between water molecules inside the VSD are highlighted in dashed lines. The hydrophobic gasket and D129 (on S1), together with the arginine residues, form two potential constriction sites breaking the continuous water wire, which in turn prevent the proton conduction through the VSD. The hydrophobic gasket is shown in surface representation for clarity.

To elucidate the mechanism of the state dependent proton conduction, we carried out MD simulations of the single cysteine mutants of the four gating residues at four major states, and analyzed continuous water wire propensity in the aqueous crevices of the VSDs (Fig. 4c, d). It has been suggested that proton conduction can be achieved if there is a continuous water wire connecting the extracellular and intracellular solutions^40,41^. R223C and R226C have a high water wire propensity at the ‘down-minus’ and ‘Down’ states, respectively. It is in good accord with our functional and computational data: the R223C mutant populates the “down-minus” state at negative potentials, resulting in steady inward proton currents as in the recordings. However, based on water wire analysis, the R226C mutant cannot conduct proton currents at the final “down-minus” state as in the R223C mutant, but can conduct in the intermediate “Down” state. This explains why only transient inward proton currents were observed in the R226C mutant mixed with the OFF gating currents, and both of which will be ceased when the VSD reaches the ending “down-minus” state.

For R232C, water molecules are able to hydrate the central hydrophobic gasket region in the ‘Up-plus’ state. However, the water wire was blocked by the salt bridge formed between R229 and D129, preventing the connection between the extracellular and intracellular solution and the steady outward proton permeation under depolarization. R229C also had a high water wire propensity at the ‘Up’ state, but its pH dependence observed experimentally was marginal. Given its large negative *V*1/2 shift (−51.70±0.73 mV compared with 53.73±0.46 mV of WT) and its faster kinetics compared with R226C, it is possible that this mutant exhibits very short ‘Up’ state dwell times (Extended Data Figs. 6a, 8a, b). In addition, the maximum net OFF charge of R229C is much smaller than others, the effect of noise and protein run-down may also weaken the pH effect (Extended Data Fig. 8c). These results further suggested that the ‘down-minus’ state is the end state of the downward movement, and the occupancy of one gating residue at the hydrophobic gasket site and the salt bridges between the gating residues and D129 are critical for preventing omega currents through the WT VSD (Extended Data Fig. 8d).

## Conclusions

In this study, we investigated the gating mechanism of Ci-VSP using the MD simulations and electrophysiological experiments to evaluate the influence of transmembrane voltage on the VSD energy landscape. We find that Ci-VSD gating cycle involves four major states, where one of the four gating charge residues occupies the center hydrophobic gasket at each state at a time. In response to voltage changes, the S4 segment undergoes a set of sequential stepwise movements: one helix turn or one ‘click’ transition at each step, which leads to the translocation of roughly one gating charge, and a total gating charge of ~3 e_0_ for the whole cycle. The ‘click’ model (Fig. 5) could be a simplified and generalized model for the movement of all the VSDs, compared with the conventional ‘helical screw’ and ‘helical sliding’ models.

**Figure 5.**
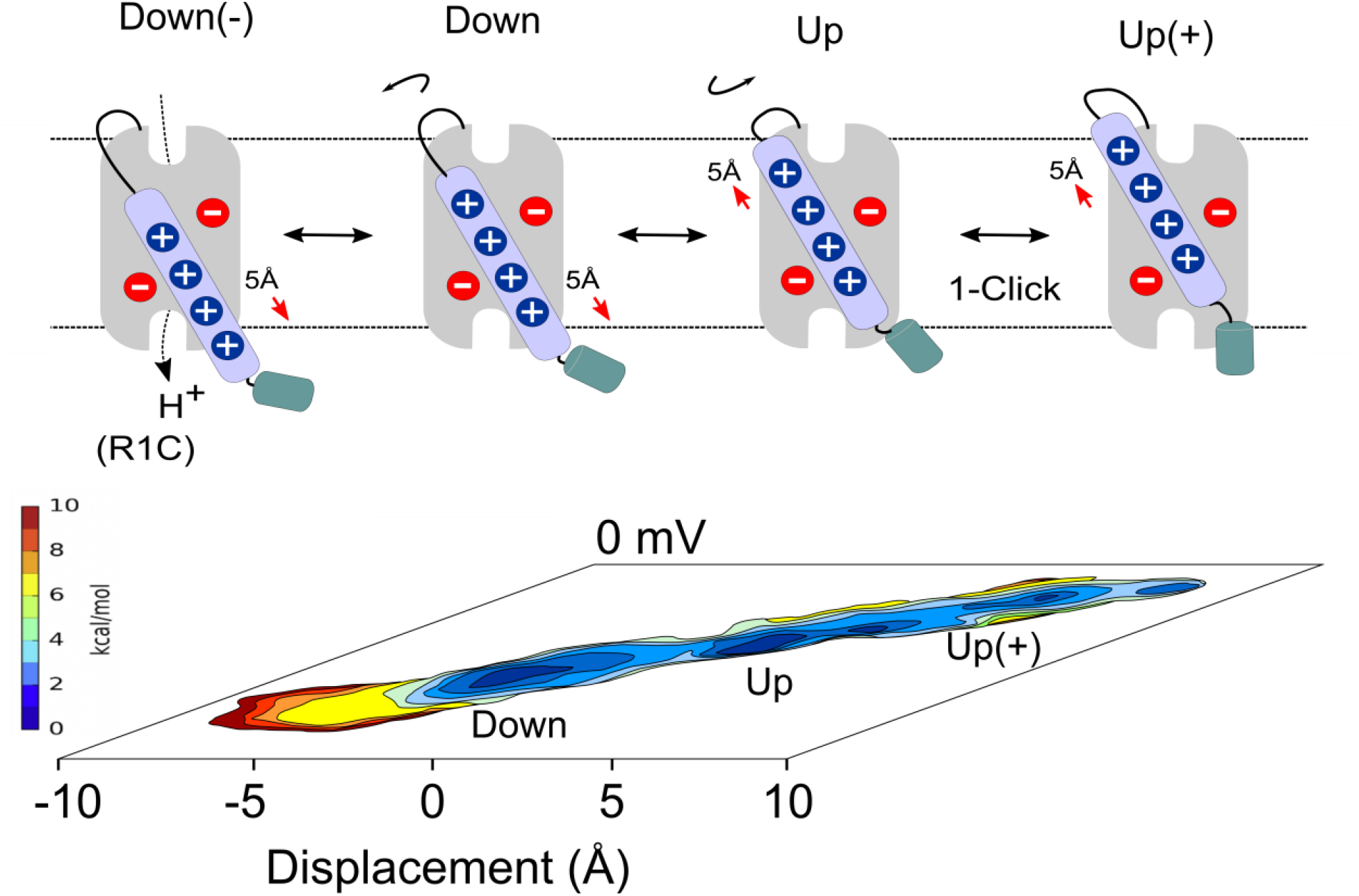
Voltage sensor gating mechanism as a series of quantized ‘clicks’. The energy landscape of a prototypical VSD (shown as a two-dimensional PMF at 0 mV) define major conformational states along the S4 movement gating cycle. In Ci-VSD, four putative conformational states, Down, Up, down-minus (Down(-)) and Up-plus (Up(+)) are related by sequential translations (of around 5 Å) and rotations to and from each adjacent state. Transmembrane voltage biases the energy landscape, triggering these transitions. Mutating the first gating charge to cysteine (R1C), leads to a proton-conductive sensor under hyperpolarizing potentials (i.e. populating the Down(-)).

Our working hypothesis is that, for hydrophobic-plug-containing sensors, S4 movement is “quantized” in the sense that S4 can only move in clicks defined by the relative position of existing countercharges. These vary according to the VSD type, explaining the wide range of voltage dependences among voltage sensors^26–29^. These clicks would be defined by either simple major S4 axis translations, rotations or secondary structure (alpha or 3_10_) helical transitions. There are, however two important caveats. First, this picture does not consider that the field might be so sharply focused that each click could effectively move more than one charge equivalent; second, it assumes that the field is constantly focused during gating. These two parameters remain largely unknown. The present results should lead to further evaluation of unsolved functional states in other VSDs by designing specific mutations and introducing state dependent metal bridges or disulfide bonds, and to investigate detailed mechanisms and potential treatment of channelopathies caused by missense mutations in VSDs.

## Acknowledgements

We thank the Francisco Bezanilla laboratory at the University of Chicago for providing oocytes and extended discussions on Ci-VSD cut-open oocyte experiments; Dr. David Medovoy for starting the free energy calculations; Dr. Giacomo Fiorin and Dr. Jérôme Hénin for advice on the usage of the Colvars module. We thank Drs. Qufei Li, Tian Li, Michael David Clark, Carlos A.Z. Bassetto, Jr., Francisco Bezanilla, and the members of the Perozo lab and the Roux lab for helpful advice and discussions. R.S. specially thanks Dr. João Carvalho-de-Souza for cut-open voltage clamp training and Ms. Wieslawa Milewski for molecular biology training. Computer resources came from an allocation on the Beagle supercomputer at the University of Chicago, and an allocation on the Blue Waters supercomputer at the National Center for Supercomputing Applications at the University of Illinois at Urbana-Champaign. This work was supported by the National Institutes of Health grants GM057846 (EP) and GM062342 (BR).

## Author contributions

R.S., B.R., E.P. designed the whole study and analyzed the data. R.S. performed the MD simulations with assistance from Y.M., and R.S. performed electrophysiological experiments. R.S. wrote the initial manuscript. R.S., Y.M., B.R. and E.P. revised the manuscript.

## Methods

### Simulation systems

The crystal structures of the “Up” (PDB: 4G7V) and “Down” (PDB: 4G80) states Ci-VSD^12^ were used as templates to build the “Up-plus” and “down-minus” states homology models. The S4 helix of the “Up-plus” and “down-minus” states VSD was shifted 3 residues upward and 6 residues downward, respectively, compared with the “Up” state Ci-VSD in the sequence alignment step of homology modeling. Then, the program MODELLER^42^ was used for the structure building of the homology models. For the purpose of consistency, the four models have the same number of residues: 106-239, with the C-terminal end of the S4 helix being modeled as a continuous helix (Extended Data Fig. 1a). The R217E mutation in the “Up” state crystal structure was reversed and its missing residues: 237-239 at the C-terminal end were rebuilt.

The “Up” state Ci-VSD was embedded into a palmitoyl oleyl phosphatidylcholine (POPC) lipid bilayer fully solvated in 0.1 M NaCl using the program VMD^43^. Titratable residues were assigned their default protonation state at pH 7. The orientation and relative positive of the protein inside the membrane was adjusted according to the prediction from the orientations of proteins in membranes (OPM) database^44^ The protein was substituted into one of the other three states Ci-VSD to construct the other three systems. The resulting four systems were electrically neutral and consisted of exactly the some components with a total number of atoms of 56,582 (Extended Data Fig. 1b).

### Molecular dynamics simulations

The systems underwent 1,000 steps of initial energy minimization followed by 30 ns and 60 ns equilibration runs for the “Up” and “Down” systems and the “Up-plus” and “down-minus” systems, respectively. The homology model structures were equilibrated 30 ns longer for better relaxation of the systems. During the equilibration runs, positions of the protein heavy atoms were harmonically restrained with the force constant being gradually decreased^45^. After the equilibration run, 20 ns production run was carried out for each of the four systems (Extended Data Fig. 1c).

### Targeted molecular dynamics simulations

Targeted molecular dynamics (TMD) simulations were performed to generate initial transition pathways between two adjacent states (Extended Data Fig. 1d). The centroids of the structures of the four systems from the last 10 ns production runs (*n* = 5,000) were used as starting and targeting configurations for the upward and downward TMD simulations. The Cartesian coordinates of the non-hydrogen atoms of the four transmembrane helices (residues: 116-138, 144-172, 181-202 and 216-236) were used in hierarchical cluster analysis to get the centroids^46^. All the heavy atoms of the protein were chosen as targeted atoms. Each TMD simulation lasted 10 ns with a spring constant of 200 kcal/mol/Å^2^ following 1,000 steps of energy minimization.

### String method with swarms of trajectories

The string method with swarms of trajectories with MD simulations was used to characterize and relax the transition pathways derived from the TMD simulations. Six strings connecting the four centroids of the structures from MD simulations were constructed and evolved for 50 iterations (Extended Data Fig. 1e). Each string contained a chain of 12 images or states. The initial configurations for the intermediate images were taken from the TMD simulations. The Cartesian coordinates of the C_α_ atoms of S4 residues: 216-235, and one representative side-chain atom of a group of key residues: the CG atoms of D129, D136, D164 and D186; the CD atom of E183; the CZ atoms of R119, R168, R217, R223, R226, R229 and R232; the OH atoms of Y200 and Y235; the OG atom of S158; the OG1 atom of T197, were selected as collective variables (Extended Data Fig. 1e). During the iteration, 20 short MD simulations (5 ps/trajectory) with different initial velocities were launched for each of the 10 intermediate images of each string. Protein structures from the last frame of the 20 trajectories of each image were then aligned, averaged and reparametrized. The resulting structure was used as the starting configuration of the image for the next iteration by a 150 ps equilibration run with collective variables being harmonically restrained (with a gradually increased force constant up to 20 kcal/mol/Å^2^) following an energy minimization of 1,000 steps^47^.

### Self-learning adaptive umbrella sampling

We used the umbrella sampling method^48^ to calculate the free energy landscape or potential of mean force (PMF) for the conformational change of Ci-VSD during the gating process. Two reaction coordinates, defined using the collective variables interface (Colvars) module^49^ in NAMD, were employed to describe the translocation and rotation of the S4 helix: *distanceZ*, projection of the distance vector between the center of mass of the C_α_ atoms of residues: 217-233 and a reference point (the center of mass of the C_α_ atoms of residues: 217-233 in the “Up” state crystal structure) along a constant vector (the principle axis of the C_α_ atoms of residues: 217-233 in the “Up” state crystal structure); and *spinAngle*, angle of the spin rotation of the C_α_ atoms of residues: 217-233 in reference to the positions of these atoms in the “Up” state crystal structure, along and around its axial axis, respectively.

To improve the efficiency of the umbrella sampling, we used the self-learning adaptive umbrella sampling approach^50^, which can explore the free energy landscape of a multi-dimensional space and find the most relevant and worth sampling regions automatically. It helps us put our efforts and computing resources on the most important subspace instead of the irrelevant high free energy regions, thus decreasing the total number of umbrella sampling windows significantly.

Snapshots from the last 10 ns production MD runs (*n* = 38) and the last ten iterations of the string method (*n* = 74) were used as starting configurations for the first cycle of the self-learning adaptive umbrella sampling simulations (Extended Data Fig. 1f). The umbrella sampling widow increments for the two reaction coordinates were 0.5 Å and 10 °, and the force constants for the two restraints were 5.0 kcal/mol/Å^2^ and 0.02 kcal/mol/deg^2^, respectively. To prevent global drifting of the protein, the projection of the center of mass of the C_α_ atoms of the S1-S3 helices (residues: 116-138, 144-172 and 181-202) in the *xy* plane and the projection of the center of mass of the phosphate atoms of lipids along the *z* axis were also restrained with a force constant of 5.0 kcal/mol/Å^2^. A free energy value of 10 kcal/mol was used as the threshold, and a total of seven cycles of 198 windows were generated. For each umbrella sampling window, a 20 ns production run was carried out following a 5 ns equilibration run with the root-mean-square deviation of the C_α_ atoms of the four transmembrane helices with respect to the starting configuration (with a force constant of 50.0 kcal/mol/Å^2^) and the orientation of the C_α_ atoms of the first three helices in reference to that in the “Up” state crystal structure (with a force constant of 10.0 kcal/mol) being also restrained. The weighted histogram analysis method (WHAM) was used to combine the sampling data and calculate the PMF^51^. The WHAM code was obtained by courtesy of Dr. Sunhwan Jo.

### Hamiltonian-replica exchange molecular dynamics (H-REMD) umbrella sampling at different membrane potentials

The landscape or the subspace explored by the self-learning adaptive umbrella sampling simulations was used in the H-REMD umbrella sampling simulations. Two one-dimensional ordered lists of umbrella sampling windows were generated to assign exchanging partners for each replica (Extended Data Fig. 1g). To keep the Euclidean distance between two exchanging replicas as short as possible^52^, we used 191 out of 198 windows derived from the self-learning adaptive umbrella sampling simulations. For the H-REMD simulations at 0 mV (no membrane potential exist), snapshots of the last frame of the selected windows from the self-learning adaptive umbrella sampling simulations were used as starting configurations directly. The reaction coordinates and restraints were the same as those in the self-learning adaptive umbrella sampling simulations. The simulations were carried out for 30 ns with an exchange attempt rate of 1 ps^−1^. For the H-REMD simulations at ±150 mV, the 191 selected self-learning adaptive umbrella sampling windows were re-simulated using the same protocol under the assigned membrane potential. The resulting configurations were used as starting structures for the 30 ns H-REMD simulations. The membrane potential was applied using the external electric field module of NAMD in which the electric field E = V/Lz, where V is the voltage and Lz is the length of the box in the z direction (scaled by the cell basis vectors)^53^. The PMFs for the H-REMD umbrella sampling simulations were also calculated via the WHAM method. The minimum free energy path connecting the local free energy minima was calculated using the zero temperature string method^54^. The average displacement charge for each sampling window was calculated as *Q*_d_ = 〈∑*_i_ q_i_z_i_*〉, using the partial charge *q_i_* and unwrapped *z_i_* coordinate of all atoms of the system from the last 5 ns trajectories of the H-REMD simulations^17,53^.

### Molecular dynamics simulations of the “Up-plus” state Ci-VSD with countercharge mutations

A double mutation (D129S/A154E) model of Ci-VSD was constructed using the program VMD based on the “Up-plus” state centroid structure from the MD simulation (Extended Data Fig. 2d). Two sets of MD simulations were performed to display the effect of the double mutation on the secondary structure conformation of the S4 helix. In one of the simulation, the distances between the atoms R226:CZ-E154:CD and R223:CZ-D136:CG were harmonically restrained (centered at 5 Å with a force constant of 5.0 kcal/mol/Å^2^) in addition to the positional restraints on the backbone atoms of the S1-S3 helices (a force constant of 1.0 kcal/mol/Å^2^) in the initial 10 ns equilibration run, followed by a 100 ns production run without any restraints. In the other simulation, the same simulation protocol was used except that the distance restraints between the two atom pairs were not implemented in the equilibration run.

### Molecular dynamics simulations of the Y200C and V220C mutants

The Y200C mutation was carried out on the centroid structures of the four systems at different states using VMD. The newly introduced cysteine residue was modeled in a deprotonated state^33^ carrying a net charge of −1 e. A Cd^2+^ ion was also added into each of the four systems (Extended Data Fig. 5d). Following 5,000 steps of energy minimization, a 5 ns equilibration run and a subsequent 10 ns production run were performed for each system. During the equilibration run, the positions of the backbone atoms of the protein and the Cd^2+^ ion initially at the center of C200:SG and D136:CG were harmonically restrained, with a force constant of 5.0 and 1.0 kcal/mol/Å^2^, respectively.

The V220C mutant was constructed on the centroid structure of the “Up-plus” state Ci-VSD using VMD. The modeling and simulation protocols were the same as the Y200C mutant, except that the Cd^2+^ was inserted between the side-chain of V200C and the phosphate atom of a nearby lipid molecule.

### Restrained molecular dynamics simulations with an explicit Cd^2+^ bridge

To explicitly simulate the Cd^2+^ bridges and assess the corresponding state they are associated with^32^, we constructed 11 double mutants of Ci-VSD in all of the four states using the centroid structures from the MD simulations with the program VMD: R217C/T197C, V220C/T197C, V220C/L137C, R223C/T197C, R226C/T197C, D213C/L137C, R226C/L137C, R217C/M133C, R226C/M133C, R226C/A154E and R229C/A154E. In addition, we constructed two more double mutants that incapable of forming Cd^2+^ bridges in our electrophysiology experiments to study the underlying molecular mechanism: R232C/A154E and D213C/M133C.

Each double mutant system was simulated for 20 ns following 5,000 steps energy minimization. The distance distribution between the Cβ atoms of the two mutated residues was calculated and analyzed together with the MD trajectory. The states in which the Cd^2+^ bridge could form were selected for the restrained MD simulation. Snapshots of the last frame of the selected trajectories were chosen to construct starting configurations for the restrained MD simulations by deprotonating the introduced cysteine residue(s) and inserting a Cd^2+^ ion between the two mutated residues using the program VMD. Each restrained MD simulation was run for 20 ns following 5,000 steps energy minimization, with the geometry of the Cd^2+^ bridge being harmonically restrained^32,33^.

### Hydration of the crevices of Ci-VSD

Single cysteine mutation of the four gating-charge residues on S4: R223C, R226C, R229C and R232C at different states were built based on the centroid structures from the previous MD simulations using the program VMD. Each system was run for 50 ns following 5,000 steps energy minimization. The breadth-first algorithm was used to find a continuous hydrogen-bonded water wire connecting the extracellular and intracellular sides of the VSD^40^. Two water molecules were considered to form a hydrogen bond when the distance between the two oxygen atoms was shorter than 3.5 Å and the angle between the donor oxygen atom, the shared hydrogen atom and the acceptor oxygen atom was larger than 120 °.

The latterly constructed systems with mutations were all neutralized, by changing the ion composition in the bulky water, if the net charge of the systems were not zero. All MD simulations were performed using NAMD^55^ with a time step of 2 fs. The CHARMM36 force field^56,57^ including the CMAP correction was used for the protein, lipids and ions, and the TIP3P model^58^ for water. The Langevin dynamics and the Nose-Hoover Langevin piston method^59,60^ were employed to control the temperature at 300 K and the pressure at 1 atm, respectively. The long range electrostatic interactions were calculated using the particle mesh Ewald (PME) method^61^ with a grid density of at least 1/Å^3^. A smoothing switch function was applied for the van der Waals interactions starting from the distance of 10 Å and with a cutoff of 12 Å.

### Molecular biology

The catalytically inactive mutant of Ci-VSP (C363S)^4^ was used as background for all the mutations and gating currents recordings, hereafter referred to as the “wild type” (“WT”) Ci-VSP. The cDNA of Ci-VSP-C363S was cloned into the pSP64T vector. All point mutations were generated using site-directed mutagenesis and confirmed by further sequencing. For all single and double cysteine mutants, the native cysteine on the voltage-sensing domain was also mutated to a serine (C159S). The plasmids containing WT or mutant Ci-VSP were linearized by restriction enzyme XbaI. cRNA was transcribed *in vitro* using the mMessage mMachine SP6 transcription kit (Ambion, Invitrogen) and diluted in RNase-free water in a concentration of approximately 1 μg/μL.

*Xenopus laevis* oocytes were surgically harvested and defolliculated using collagenase solution with bovine serum albumin (BSA). Each freshly isolated oocyte was injected with 50 nL of cRNA. Injected oocytes were incubated for 16–24 hours at 18 °C in standard oocyte saline (SOS) solution containing the following components: 100 mM NaCl, 5 mM KCl, 2 mM CaCl_2_, 1mM MgCl_2_, 10 mM HEPES at pH 7.4, and 50 μg/ml gentamycin. For the various cysteine mutants, 0.5 mM dithiothreitol (DTT) was added into the incubation solution to prevent formation of disulfide bond^10^ and chelate heavy metal contaminants from water and/or chemicals^62,63^.

### Electrophysiological recordings and data analysis

Gating currents were recorded at room temperature (20–23 °C) on a cut-open oocyte voltage-clamp setup^64,65^. Raw currents were filtered at 10 kHZ with a low pass four-pole Bessel filter within the CA-1 amplifier (Dagan Corporation; Minneapolis, MN). The ON and OFF currents were evoked using the following voltage protocol: in brief, from a holding potential of −60 mV, a 200 ms conditioning prepulse to −100 mV was applied to maximally deactivate Ci-VSP. Then, 250 ms depolarization voltage steps from 160 mV to −100 mV in −10 mV decrements were applied to initiate the ON currents, followed by a 350 ms repolarization potential to −100 mV to measure the OFF currents. Due to the presence of non-negligible endogenous outward currents at potentials more positive than 40 mV^66^, with subtle inward currents upon returning to very negative potentials (Extended Data Fig. 4a), we used the net OFF charge to represent the total charge movement *Q* which was calculated by time integration of the OFF gating current. For mutants whose gating charge starts moving at more negative potentials, the prepulse and repolarization potentials and the minimum test pulse were changed correspondingly to more negative potentials (e.g., R229C to −140 mV). For mutants involving the R223C mutation which conduct omega currents at polarization states^35,36^, preceded by a 200 ms prepulse to 0 mV, 800 ms voltage steps from 20 mV to −150 mV were applied to evoke the currents followed by a 400 ms voltage step to 0 mV, and the holding potential was 40 mV. The steady-state current amplitudes *I* were used to study the functional property of the mutants. Leak and fast transient capacitive currents were subtracted offline. Data acquisition and analysis were performed using in-house software GPatch and Analysis, respectively, kindly provided by Prof. Francisco Bezanilla.

The external solution contained (in mM): 120 N-methyl-D-glucamine (NMDG), 2 Ca(OH)2, 0.5 EDTA and 10 buffering agent: HEPES for pH7.4, MES for pH6.0 and CHES for pH9.0. The internal solution contained (in mM): 120 NMDG, 2 EGTA, and 10 HEPES at pH7.4. Methanesulfonic acid was used for pH adjustment. When needed, Cd^2+^ was diluted in the external solution without EDTA at designed concentration from a 100 mM CdCl_2_ stock solution^67^. 1~2 mM DTT was freshly added to the external solution to chelate/washout the Cd^2+^ ions.

The normalized net OFF charge-voltage (*Q–V*) curves for each construct and condition (extracellular recording solution) were averaged and fitted with a Boltzmann distribution, *Q*(*V*)=1/{1+exp[*ze*_0_(*V*–*V*_1/2_)/*k*_B_*T*]}, where *z* is the apparent gating charge, *V*_1/2_ is the half activation voltage, *e*_0_ is the elementary charge, *k*_B_ is the Boltzmann constant and *T* is the absolute temperature in Kelvin. The python scipy.optimize.curve_fit procedure was used for fitting and calculating the best-fit values and standard deviations for the parameters *z* and *V*_1/2_. The OFF gating currents were fitted to a sum of two independent exponential functions, and a weighted mean time constant, *τ*_OFF_=(*A*_1_*τ*_1_+ *A*_2_*τ*_2_)/(*A*_1_+*A*_2_), was used to characterized the kinetics of deactivation, where *A*_1_, *A*_2_ and *τ*_1_, *τ*_2_ are the amplitudes and time constants for the first and second exponentials, respectively. The effective Cd^2+^ effect on the maximum normalized net OFF charge of the double mutants, with respect to the maximum net OFF charge in the absence of Cd^2+^, was calculated by subtracting the corresponding Cd^2+^ effect on each single mutant from that of the double mutant, ΔΔ*Q*_ij_/*Q*_ij, max, w/o Cd^2+^_ = |*Q*_ij_/*Q*_ij, max, w/o Cd^2+^_ – *Q*_i_/*Q*_i, max, w/o Cd^2+^_* *Q*_i_/*Q*_i, max, w/o Cd^2+^_|, with the standard deviation being estimated using the rule of linear propagation of uncertainty^68^.

