## Supplemental figures for "Mechanism of Voltage Gating in the Voltage-Sensing Phosphatase Ci-VSP"

### Extended Data Figures

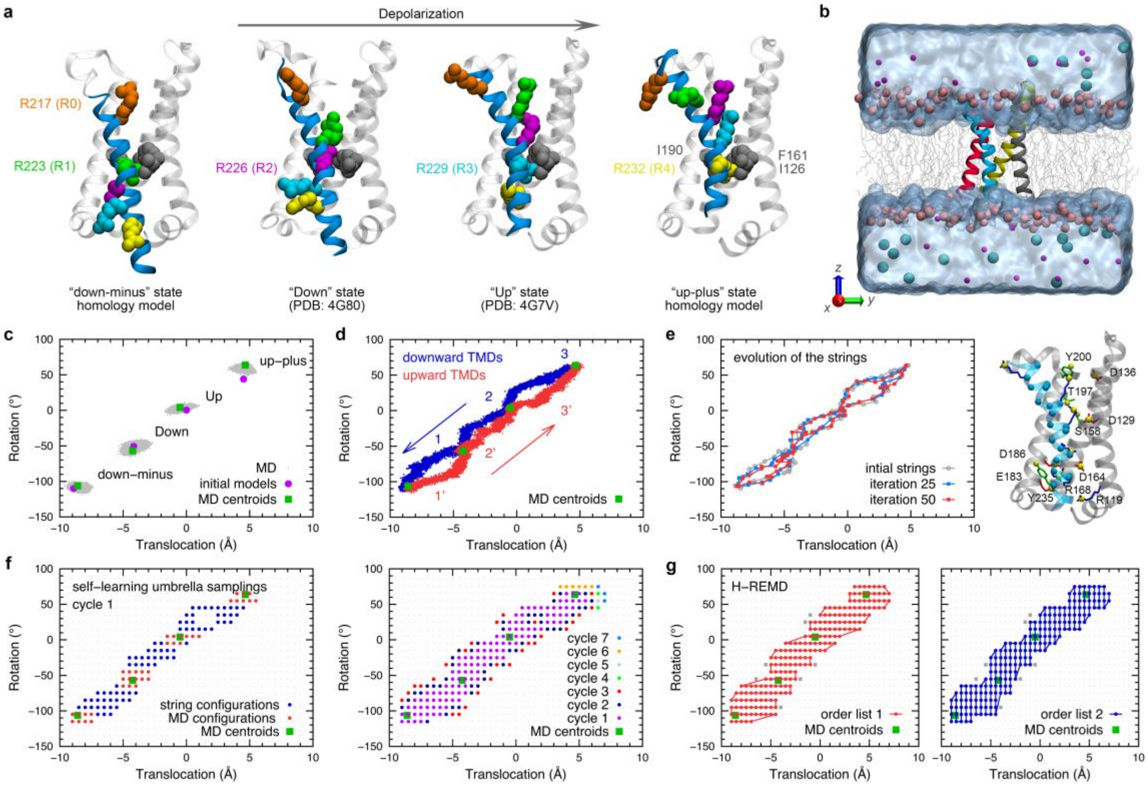


**Extended Data Figure 1. Workflow for the molecular dynamics (MD) studies of the gating process of the voltage-sensing domain (VSD). a**, Starting configurations of the VSD in the "down-minus", "Down", "Up" and "up-plus" states. The arginine residues on S4 and hydrophobic gasket residues (I126, F161 and I190) are shown in vdW representation. **b**, A MD simulation model of the VSD. The four transmembrane helices are shown in ribbon representation: S1 (gray), S2 (yellow), S3 (red) and S4 (blue); the lipids in gray line representation with the phosphate atoms in pink spheres; bulky water in surface; and ions in spheres. **c**, Projection of the VSD structures from the last 10 ns trajectory of each MD simulation ($n$ = 5,000, gray dots), the initial models (purple circle) and the centroids of the VSD structures from the last 10 ns trajectory of each MD simulation (green square) into the two reaction coordinates describing the movement of the S4 helix. **d**, Projection of the VSD structures from the targeted molecular dynamics (TMD) simulations. **e**, Left, projection of the VSD structure in each of the windows composing the string paths from iterations 0 (gray), 25 (blue) and 50 (red) into the two reaction coordinates, showing the evolution of the string paths connecting the adjacent states. Right, the 36 selected atoms whose Cartesian coordinates having been used as collective variables in the string method are highlighted in spheres. The "Up" state VSD (PDB: 4G7V) is shown in ribbon representation, with the S4 helix being colored in blue for clarity. **f**, Left, snapshots from the MD simulations (red) and the windows of the string paths (blue) were used as starting configurations for the self-learning umbrella sampling simulations. Right, seven cycles of umbrella sampling simulations were performed. The restraint centers of the umbrella sampling simulations in each cycle are shown in different colors. **g**, The two order lists used in the Hamiltonian-replica exchange molecular dynamics (H-REMD) umbrella sampling simulations. Left, the “translocation” is the fast changing reaction coordinate. Right, the “rotation” is the fast changing reaction coordinate. The centers of those self-learning umbrella sampling windows that not involved in the H-REMD simulations (7 out of 198) are shown in gray.

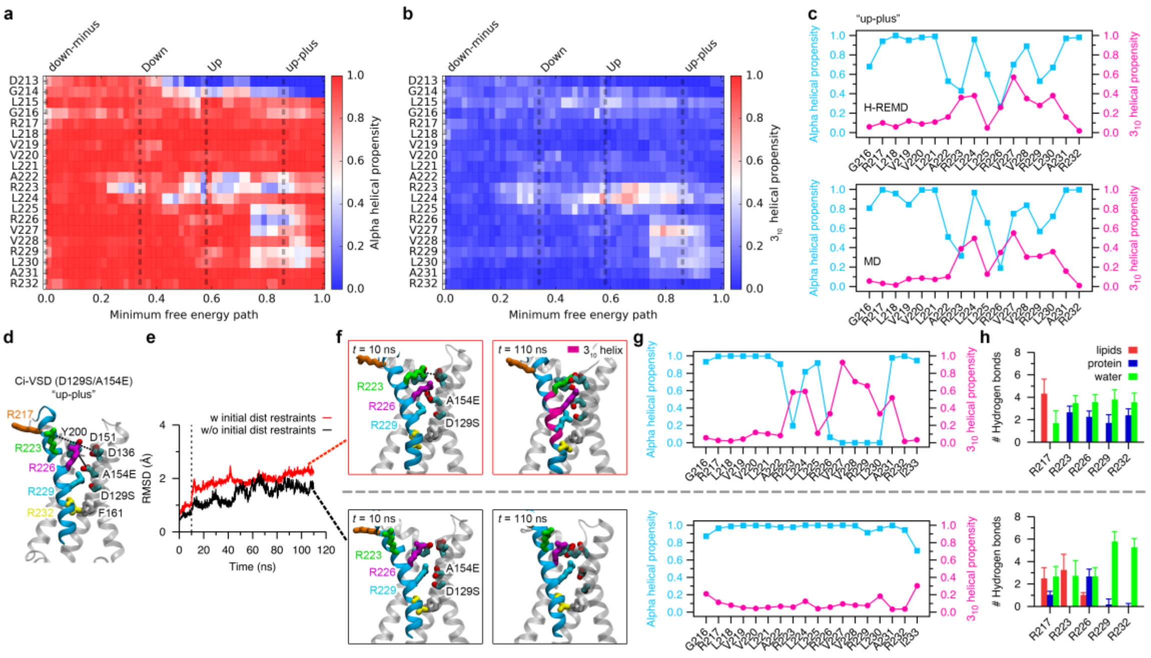


**Extended Data Figure 2. Secondary structure of the S4 helix.** **a**, Propensity of S4 residues adopting an alpha helical conformation along the minimum free energy path. **b**, Propensity of S4 residues adopting a $3_{10}$ helical conformation along the minimum free energy path. Residue *i* is classified to adopt an alpha helical or a $3_{10}$ helical conformation when it forms a hydrogen bond ($d_{O\text{-}N}$ < 3.5 Å and $\alpha_{O\text{-}H\text{-}N}$ > 120 °) to residue *i*+4 or *i*+3, respectively. In **a** and **b**, the VSD structures from the last 5 ns trajectory of the H-REMD simulation at 0 mV were used for the calculation ($n$ = 100). The vertical dashed lines represent the approximate locations of the four major states of the VSD along the minimum free energy path. **c**, The alpha helical and $3_{10}$ helical propensities for S4 residues of the "up-plus" state VSD during the H-REMD (top, data from **a** and **b**) and MD simulations (bottom). The VSD structures from the last 10 ns trajectory of the MD simulation were used for the calculation ($n$ = 1,000). **d–h**, MD simulations of the double-mutant Ci-VSD (D129S/A154E) showing perturbation of the secondary structure of S4 by locations of the countercharges and hydrogen bond interactions between the gating charges and the polar residues and lipid molecules. **d**, Starting configuration of the double mutant VSD in the "up-plus" state. **e**, RMSDs of the backbone atoms of the four transmembrane helices in two 110 ns MD simulations. The distances between atoms R226:CZ-E154:CD and R223:CZ-D136:CG, highlighted with dashed lines in **d**, were harmonically restrained (centered at 5 Å with a force constant of 2 kcal/mol/Å$^2$) at the first 10 ns of one simulation (red line). **f**, Snapshots of the VSD taken at time *t* = 10 ns and 110 ns of the two simulations with (top) and without (bottom) initial distance restraints were shown. The $3_{10}$ helical region of the S4 helix is colored magenta. **g**, The alpha helical and $3_{10}$ helical propensities for S4 residues in the two simulations. **h**, The averaged number of hydrogen bonds formed by each of the five arginine residues on S4 with lipid molecules (red), polar residues on S1-

S3 (blue) and water molecules (green) in the two simulations. Error bars denote standard deviation. The VSD structures from the last 10 ns trajectory of the simulations were used for the calculation in **g** and **h** ($n$ = 1,000).

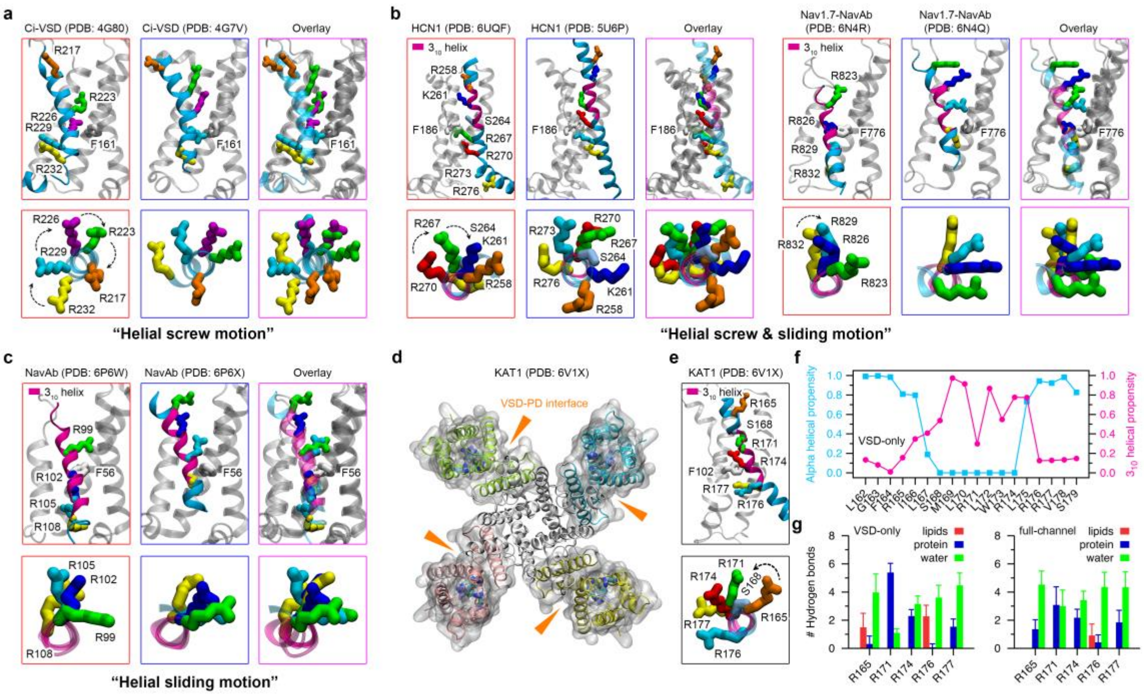


**Extended Data Figure 3. Proposed gating mechanisms in different types of voltage-sensing domains.** **a**, Crystal structures of the VSD of Ci-VSP at two different states. **b**, Cryo-EM structures of the VSD of the HCN1 channel and the Nav1.7-NavAb chimera channel at two different states. **c**, Cryo-EM structures of the VSD of the NavAb channel at two different states. **d**, Top view of the cryo-EM structure of the KAT1 channel. The S1-S5 helices from different subunits are colored differently with transparent surfaces. **e**, Close view of the VSD of KAT1. **f**, The alpha helical and $3_{10}$ helical propensities for S4 residues of KAT1. The VSD structures from the last 10 ns trajectory of the 50 ns simulation of KAT1 VSD-only system were used for the calculation (*n* = 1,000). **g**, The averaged number of hydrogen bonds formed by the arginine residues on S4 of KAT1 with lipid molecules (red), polar residues on S1-S3 (blue) and water molecules (green) in the simulations with an isolated VSD (left) and a full-channel (right). Error bars denote standard deviation. The snapshots from the last 10 ns trajectory of the 50 ns simulation of the KAT1 VSD-only system (*n* = 1,000) and the 40 - 50 ns trajectory of the 500 ns simulation of the KAT1 full-channel system (*n* = 1,000) were used for the analysis. The VSD and PD interfaces, as shown in **d**, prevent the formation of hydrogen bond interactions between the gating charges on S4 and the lipid bilayer. The dashed arrows in the top view of the S4 helices (bottom panels in **a**, **b** and **e**) represent approximate rotating motions of certain positively charged residues during the gating process. The $3_{10}$ helical parts of the helices are colored magenta. S4 residues in the alpha helical and $3_{10}$ helical conformations will undergo helical screw and helical sliding motions during gating, respectively.

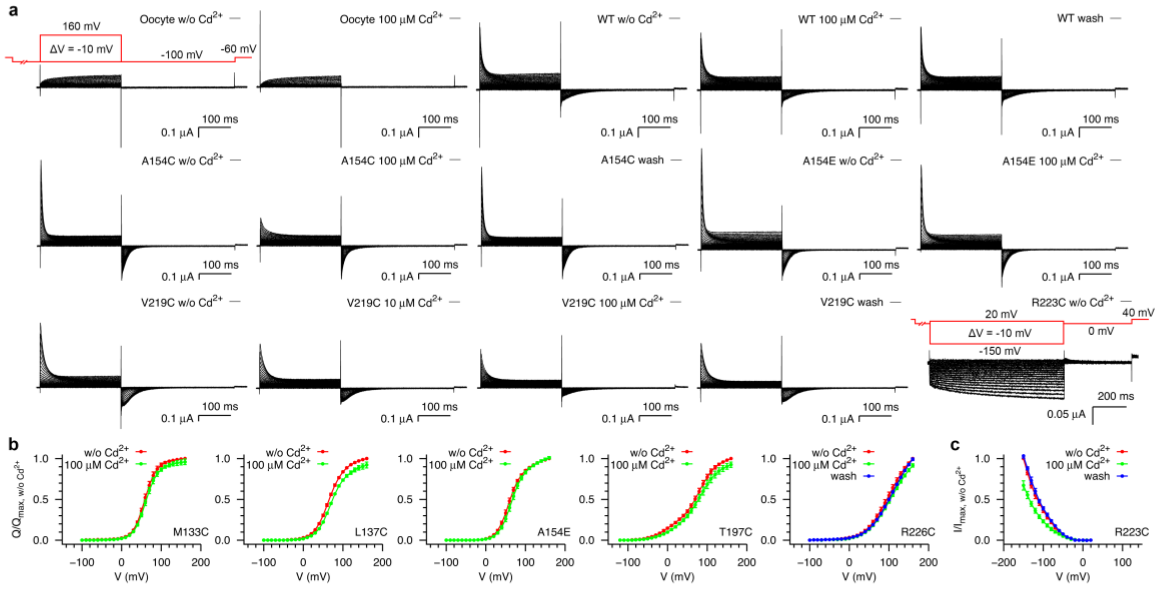


**Extended Data Figure 4. Functional characterization of wide type and different mutants of Ci-VSP in response to external $Cd^{2+}$ ions.** **a**, Representative currents for non-injected *Xenopus* oocytes and oocytes expressing wide type (WT) and single mutants of Ci-VSP without (w/o), in the presence of, and after washout of extracellular $Cd^{2+}$ ions. A different voltage protocol (inset) was used for Ci-VSP/R223C as omega currents will be elicited at negative voltages for the mutant. **b**, Representative normalized *Q*-*V* curves for single mutants of Ci-VSP. OFF gating currents were integrated to yield the net translated charge *Q*. The *Q*-*V* curve for each experiment was normalized with respect to the corresponding maximum *Q* measured in the absence of $Cd^{2+}$. **c**, Representative normalized *I*-*V* curve for Ci-VSP/R223C. After linear leak subtraction, the *I*-*V* curve for each experiment was normalized with respect to the corresponding maximum current amplitude at the end of the pulses measured in the absence of $Cd^{2+}$. Error bars denote standard deviation (M133C, $n$ = 4; L137C, $n$ = 5; A154E, $n$ = 3; T197C, $n$ = 4; Y200C, $n$ = 5; R223C, $n$ = 3).

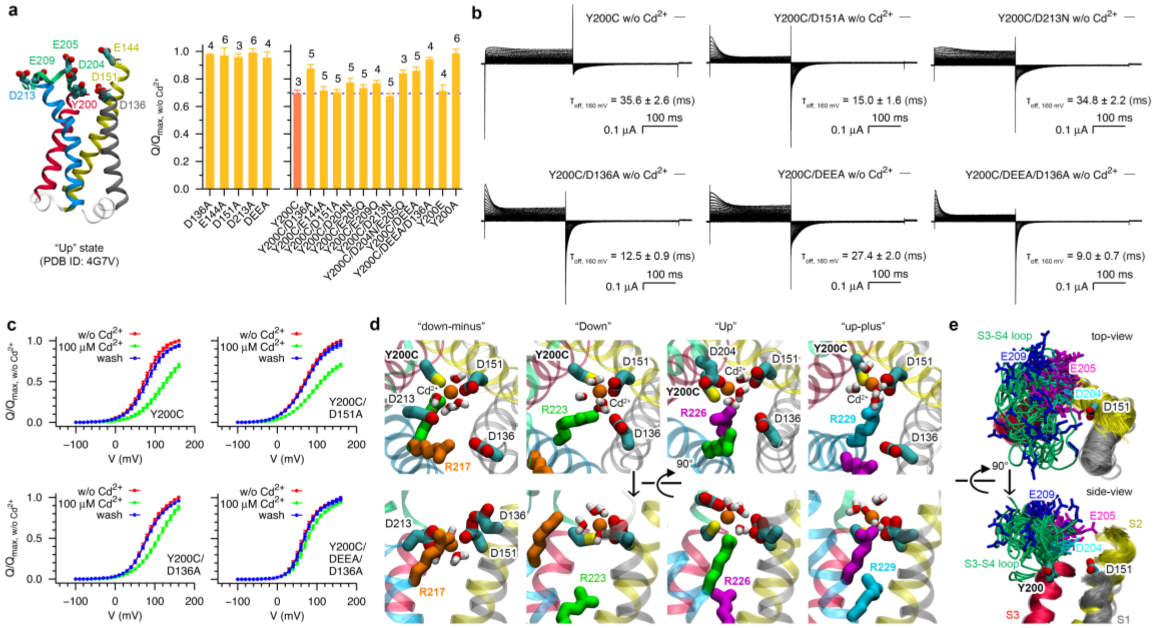


**Extended Data Figure 5. $Cd^{2+}$ effect on the Y200C mutation at the extracellular side of the S3 helix. a**, Left, crystal structure of the "Up" state VSD showing the negatively charged residues surrounding Y200. Right, the relative decreasing of the maximum net off gating charge of Y200C included mutants in response to 100 μM $Cd^{2+}$. The dashed line represents the mean value of that for the single mutant Y200C as a reference. Error bars denote standard deviation with the number of experiments being listed on the top of each of the bars. **b**, Representative currents for the mutants of Ci-VSP in the absence of extracellular $Cd^{2+}$ ions. Insets are the time constant (mean ± standard deviation) of the OFF gating current at the voltage pulse of 160 mV. The number of experiments for each construct is in **a**. **c**, Representative normalized *Q*-*V* curves. **d**, Top view (top) and side view (bottom) of snapshots from MD simulations of the VSD with the Y200C mutant at the four major states. A $Cd^{2+}$ ion was initially placed at the center of the residues Y200C and D136. Water molecules at the first solvation layer of $Cd^{2+}$ are highlighted in sticks. **e**, Overlay of snapshots ($n$ = 51) from simulation windows along the minimum free energy path of the H-REMD umbrella sampling simulation at 0 mV. The flexible S3-S4 loop and the three negatively charged residues in it are shown in tube and stick representations, respectively. The S4 helix and the S1-S2 loop are removed for clarity.

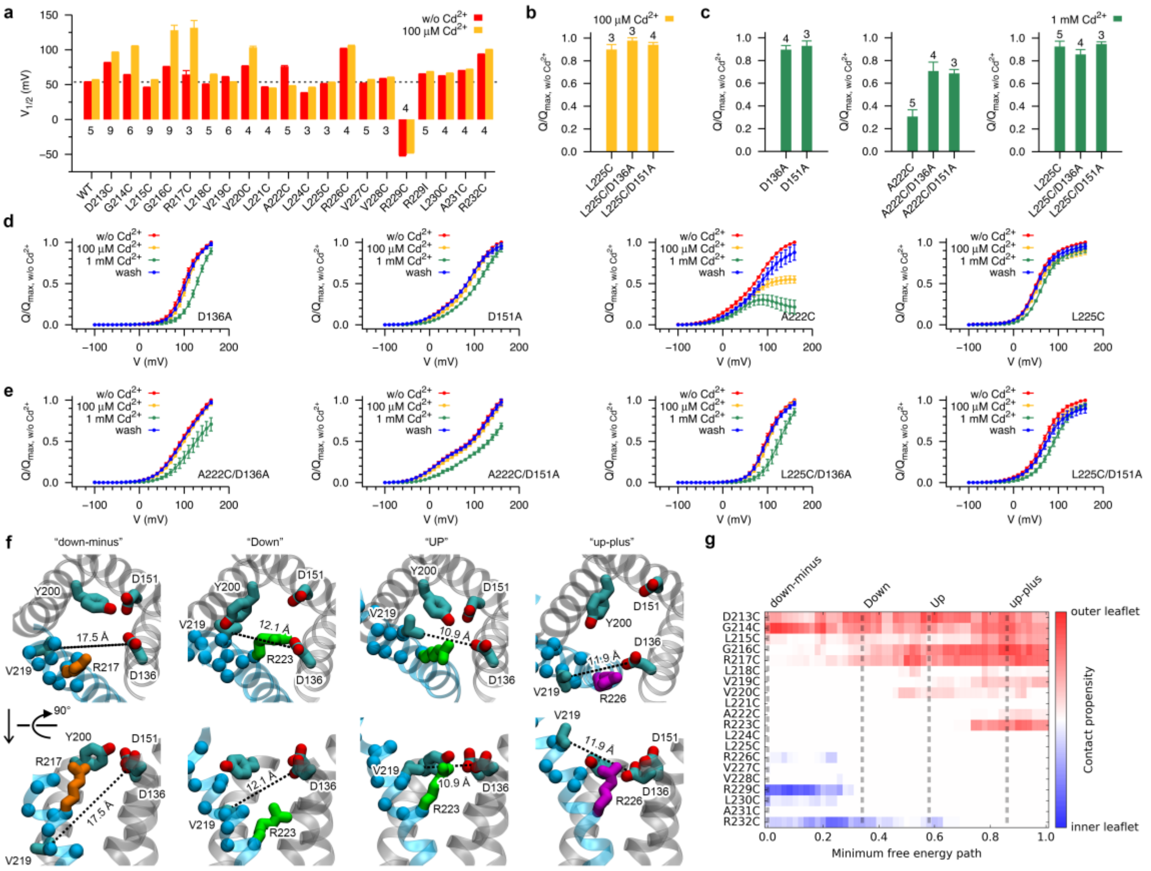

**Extended Data Figure 6. Single cysteine mutants at the extracellular side of S4 could form $Cd^{2+}$ bridges with the outer leaflet lipid. a**, Perturbation of the half-activation voltage ($V_{1/2}$) by single cysteine mutations on the S4 helix and the presence of 100 μM $Cd^{2+}$ in the external recording buffer. **b-e**, The extent of immobilization of the S4 helix is regulated by the external $Cd^{2+}$ concentration and the energetics of gating. **b**, The relative decreasing of the maximum net OFF gating charge of mutants of Ci-VSP in response to 100 μM $Cd^{2+}$. **c**, The relative decreasing of the maximum net OFF gating charge of WT and mutants of Ci-VSP in response to 1 mM $Cd^{2+}$. **d**, The normalized *Q*-*V* curves for the WT and single mutants of Ci-VSP. **e**, The normalized *Q*-*V* curves for the double mutants of Ci-VSP. **f**, Top (top) and side (bottom) view of snapshots of the VSD from MD simulations showing that cysteine mutants at the extracellular side of S4 can hardly form $Cd^{2+}$ bridges with D136. Alpha carbon atoms of residues 213-223 are shown in spheres. The distance between the two atoms V219:CB and D136:CG (connected with a dashed line) in each configuration has been labeled. **g**, Contact (or $Cd^{2+}$ bridge formation) propensity of S4 residues with the outer and inner leaflets of lipids along the minimum free energy path. The snapshots from the last 5 ns trajectories of simulation windows along the minimum free energy path of the H-REMD umbrella sampling simulation at 0 mV were used for the calculation ($n$ = 50). Residue $i$ is considered to contact to a lipid when the distance between its alpha carbon atom and any one of the nonester phosphate oxygen atoms is below 6 Å.

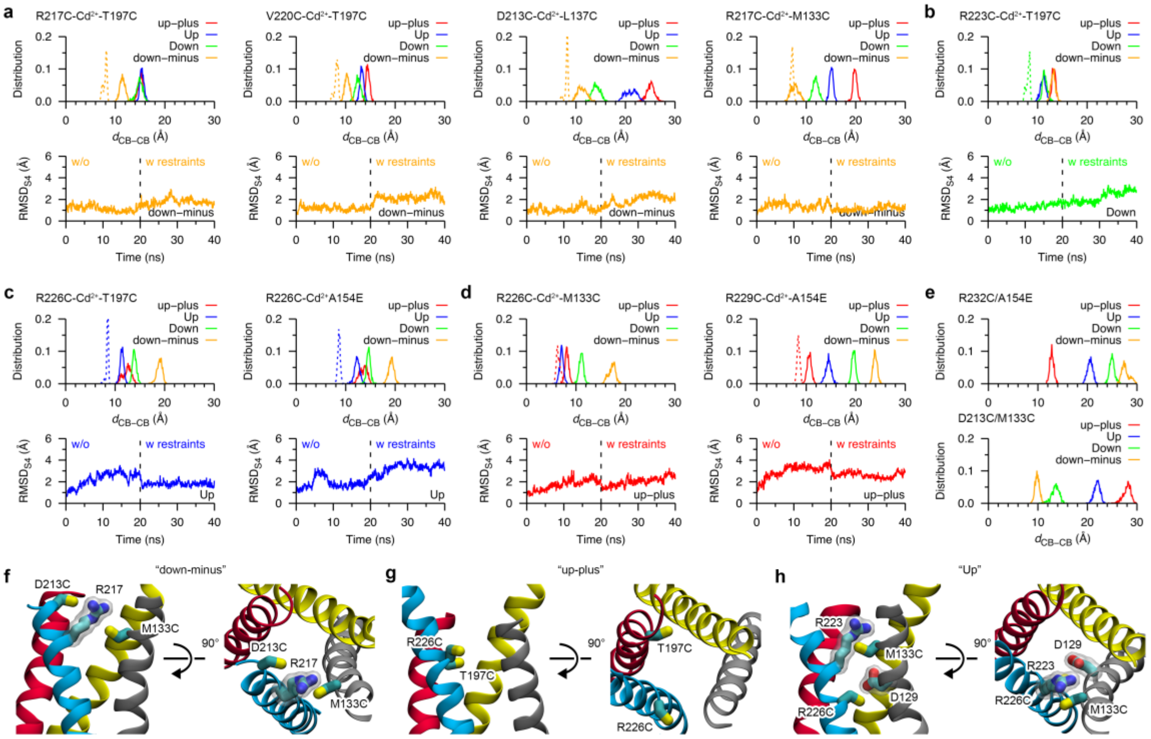

**Extended Data Figure 7. MD simulations of the double mutants of Ci-VSD with an explicit $Cd^{2+}$ bridge. a-d**, Top, the distance distribution of the beta carbon atoms of the two mutated residues in each of the 20 ns MD simulations of the Ci-VSD mutants in the "up-plus" (red), "Up" (blue), "Down" (green) and "down-minus" (orange) states (solid line), and in the 20 ns restrained MD simulation of the Ci-VSD mutants in a specific state (the same color scheme) with harmonic restraints on the $Cd^{2+}$ bridge geometry (dashed line). Bottom, the time series of the backbone RMSD of the S4 helix in the MD (black line) and restrained MD (color line) simulations of the Ci-VSD mutants in the "down-minus" (**a**), "Down" (**b**), "Up" (**c**) and "up-plus" (**d**) states. **e**, The distance distribution of the beta carbon atoms of the two mutated residues in each of the 20 ns MD simulations of the two double mutants of Ci-VSD (R232C/A514E and D213C/M133C) in different states. These two double mutants cannot form $Cd^{2+}$ bridges in experiments. Snapshots from the last 10 ns trajectory of each simulation were used for the calculation of the distance distributions ($n$ = 1,000). **f-h**, Distance is not the only factor preventing the formation of a $Cd^{2+}$ bridge between two cysteine residues. Intruding of another residue between the two cysteine residues (**f**), impropriate orientation of the side-chain of the two cysteine residues facing away from each other (**g**), and blockage of $Cd^{2+}$ from getting into the middle of the two cysteine residues by other residues (**h**) will all prohibit the formation of a $Cd^{2+}$ bridge even thorough the two cysteine residues are in an appropriate distance.

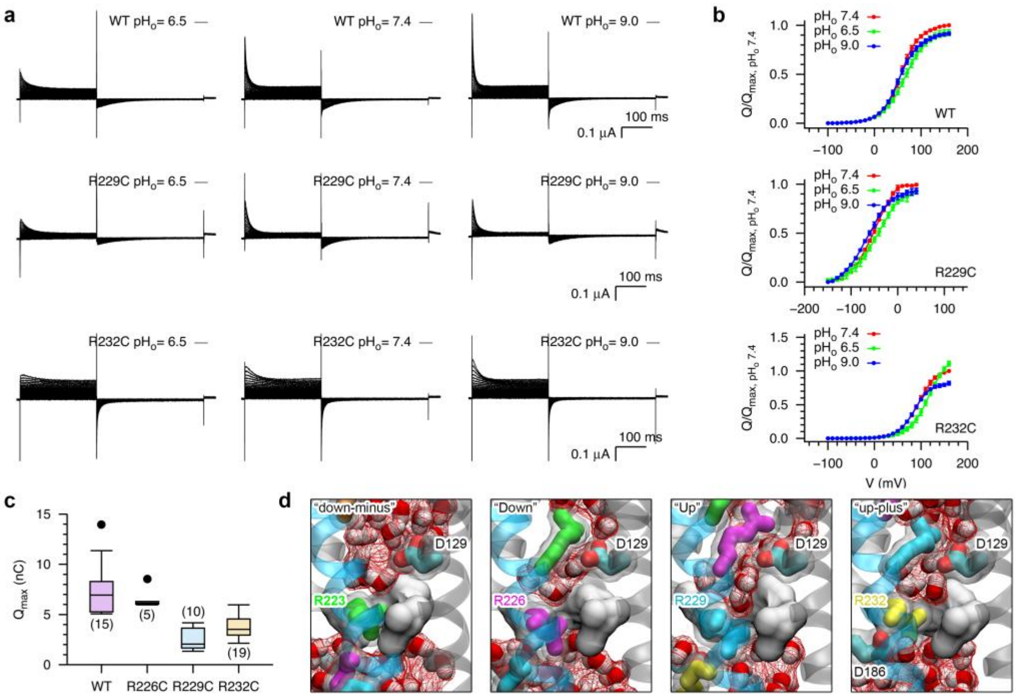


**Extended Data Figure 8. Determinants preventing the proton leaking through the voltage-sensing domain of Ci-VSP.** **a**, Representative currents for WT, R229C and R232C mutants of Ci-VSP at different pH values ($pH_o$) of the external recording buffer. **b**, Normalized *Q*-*V* curves for WT, R229C and R232C mutants of Ci-VSP. The *Q*-*V* curve for each experiment was normalized with respect to the corresponding maximum *Q* measured at $pH_o$ 7.4. Error bars are standard deviation (WT, $n$ = 7, 5, 5 for $pH_o$ 7.4, $pH_o$ 6.5, $pH_o$ 9.0, respectively; R229C, $n$ = 4, 5, 6 for $pH_o$ 7.4, $pH_o$ 6.5, $pH_o$ 9.0, respectively; R232C, $n$ = 4, 9, 6 for $pH_o$ 7.4, $pH_o$ 6.5, $pH_o$ 9.0, respectively). **c**, Box plot of the maximum net OFF charges of WT, R226C, R229C and R232C mutants of Ci-VSP. The R229C mutant has a smaller $Q_{max}$ compared with the others. The number of experiments for each construct is listed inside the parentheses. **d**, Snapshots of the WT VSD in the "down-minus", "Down", "Up" and "up-plus" states. Water molecules inside the VSD are shown in sticks and wireframe surface representation. The hydrophobic gasket is shown in solid surface representation. The gating charge (R223, R226, R229 or R232) and the surrounding hydrophobic gasket residues form one constriction site, and the countercharge D129 on S1 form the other constriction site by forming salt bridges with the gating charges, to prohibit the formation of a continuous water wire connecting the two sides of the VSD, thus prevent the proton leaking during the gating process.
